# The Unexpected Role of GCN2 Kinase Activation in Mediating Pulmonary Arterial Hypertension

**DOI:** 10.1101/2023.09.05.556450

**Authors:** Maggie M. Zhu, Jingbo Dai, Zhiyu Dai, Yi Peng, You-Yang Zhao

## Abstract

**Background:** Pulmonary arterial hypertension (PAH) is characterized by progressive increase of pulmonary vascular resistance and remodeling that result in right heart hypertrophy and failure. Published studies show that recessive mutations of *EIF2AK4* gene (encoding GCN2, General control nonderepressibe 2 kinase) are linked to heritable pulmonary veno-occlusive disease (PVOD) in patients and *EIF2AK4* mutations were also found in PAH patients although very rare. However, the role of GCN2 kinase in the pathogenesis of PAH remains unclear.

**Methods:** *Eif2ak4^-/-^* mice with genetic disruption of the kinase domain and GCN2 kinase inhibitor A-92 were employed in animal models of PH including chronic hypoxia-exposed mice and monocrotaline-challenged rats. Human lung endothelial cells (HLMVECs) were used for mechanistic studies. Endothelium-targeted nanoparticles were employed to deliver plasmid DNA to adult mice to knockout *Eif2ak4* or overexpress *Endothelin-1 (Edn1)* selectively in ECs.

**Results:** Here we show that loss of GCN2 induced neither spontaneous PVOD nor PH in *Eif2ak4^-/-^* mice but inhibited hypoxia-induced PH evident by reduced right ventricular systolic pressure, right ventricle hypertrophy and pulmonary vascular remodeling. RNA sequencing analysis suggested Edn1 as the downstream target of GCN2. In cultured HLMVECs, GCN2 was phosphorylated and activated in response to hypoxia, mediating hypoxia-induced Edn1 expression via HIF-2α. Restored *Edn1* expression in ECs in *Gcn2*-deficient mice reversed the reduced phenotype of hypoxia-induced PH. Furthermore, loss of endothelial *Eif2ak4* in mice attenuated hypoxia-induced PH. Monocrotaline-induced PH and pulmonary vascular remodeling in rats were inhibited by GCN2 inhibitor A-92 treatment. The clinical relevance of the observation was validated by GCN2 hyperphosphorylation indicative of activation in ECs of pulmonary vascular lesions of PAH patients.

**Conclusion:** These studies demonstrate that GCN2 activation by hypoxia mediates pulmonary vascular remodeling and PAH through Edn1. Thus, targeting GCN2 signaling is a promising therapeutic strategy for treatment of PAH in patients without *EIF2AK4* loss of function mutations.

## Introduction

Pulmonary hypertension (PH) is defined by a resting mean pulmonary arterial pressure greater than 20 mmHg.^1^ Although there are many causes of PH, the development of PH is almost always associated with exacerbating symptoms and increased mortality, regardless of the underlying disease.^2^ Based on pathophysiological mechanisms, clinical presentation, hemodynamic characteristics and therapeutic management, PH is clinically classified into five groups.^1^ Pulmonary arterial hypertension (PAH) (group 1) is characterized by a progressive increase in pulmonary vascular remodeling and pulmonary vascular resistance, resulting in right ventricle hypertrophy and, ultimately, right heart failure.^3–5^ Over the past few decades, progress has been made for the treatment of PAH and substantially improved the prognosis of the patients with PAH.^6–9^ However, currently approved PAH medications are mainly focusing on reducing pulmonary vascular constriction and only provide partial symptomatic relief.^10^ Due to the poor understanding of the molecular mechanisms of pulmonary vascular cell dysfunction leading to the progressive vascular remodeling, there is a lack of treatment directly targeting pulmonary vascular remodeling, and thus the mortality rate of PAH remains unacceptably high, which is still 50% at 5 years after diagnosis.^11^

General control nonderepressible 2 (GCN2), a serine-threonine kinase found in all eukaryotic organisms, is primarily known as a sensor of metabolic stress such as limited amino acids, glucose, or purine.^12–16^ In response to amino acid starvation, GCN2 is activated by the binding of uncharged tRNAs accumulated in amino acid-starved cells to the histidyl-tRNA synthetase (HisRS)-related domain of GCN2 ^17–19^ inducing conformational changes ^20–21^ and autophosphorylation,^22^ and subsequently phosphorylates the subunit of eukaryotic translation initiation factor 2 (EIF2α) to suppress general protein synthesis but selectively activate stress protein synthesis for adaptation to amino acid starvation.^14^ Autophosphorylation in the activation loop of GCN2, especially at yeast GCN2 amino acid Thr887 equivalent to mouse Thr898 and human Thr899, is required for GCN2 activation^22^ and it locks the kinase domain in an open active form which can bind its substrate such as EIF2α in the absence of tRNA.^21–25^ In addition to sensing nutrition deprivation, recent studies have shown that GCN2 can also be activated by several other stresses including UV irradiation,^26^ oxidative stress,^27^ and hypoxia.^28^ Apart from the role in regulating translation initiation, GCN2 has also been implicated in affecting G1 arrest and apoptosis,^28^ tumor growth,^29^ and inflammation.^30^

Recessive mutations in the *EIF2AK4* gene (encoding GCN2) and resultant reduction of GCN2 are linked to heritable pulmonary veno-occlusive disease (PVOD), a rare subgroup of severe PAH, which is characterized by intimal proliferation and fibrosis of septal veins and pre-septal venules.^31^ The PVOD subgroup has a worse prognosis than the classical PAH group and currently there are no effective treatments in addition to lung transplantation.^32^ Biallelic *EIF2AK4* mutations are found in 25% of histologically confirmed sporadic cases of PVOD.^31^ Although *EIF2AK4* mutation is rarely identified in idiopathic PAH (IPAH) patients (9 of 864 PAH patients),^33^ *EIF2AK4* mutation is also identified in some heritable PAH patients but not in family members without PAH.^34^ Most of these mutations are stop codons or insertions/deletions that disrupt gene function, indicating that loss of GCN2 function induces PVOD or PAH without PVOD in patients.

Here we sought to determine whether *Eif2ak4^-/-^* (KO) mice with disruption of the kinase domain develop spontaneous PVOD and the role of GCN2 signaling in pulmonary vascular remodeling and PAH development. To our surprise, The KO mice didn’t develop spontaneous PVOD and PH. Thus, we challenged the mice with chronic hypoxia. Unexpectedly, loss of Gcn2 in the KO mice attenuated PH, which was caused by reduced Edn1 expression. Furthermore, CRISPR/Cas9-mediated knockout of endothelial *Eif2ak4* in adult mice attenuated hypoxia-induced PH. GCN2 kinase inhibitor A-92 treatment also inhibited monocrotaline-induced PH in rats. In IPAH patients, we observed prominent GCN2 phosphorylation at Thr 899, i.e. GCN2 activation in pulmonary vascular ECs with little changes of its mRNA and protein expression levels compared to lungs of normal donors. To the best of our knowledge, our studies for the first time demonstrate a detrimental role of GCN2 activation in promoting pulmonary vascular remodeling and PAH development. Thus, targeting GCN2 signaling may represent a novel therapeutic approach for treatment of PAH patients without *EIF2AK4* recessive mutations.

## Methods

### Data availability

Detailed Methods and Major Resources Table are available in the Online Supplemental Material. RNA sequencing data were deposited at NCBI Sequencing Read Archive database (SRA accession#PRJNA994888). The data that support the findings of this study, experimental materials, and analytic methods are available from the corresponding author, upon reasonable request.

### Statistical analysis

Prism 8 (GraphPad Software, Inc.) was used for statistical analysis. Two-group comparisons were analyzed by the unpaired two-tailed *t* test for normal distribution or Mann-Whitney *U* test for non-normal distribution. Multiple group comparisons were analyzed by one-way ANOVA followed by Sidak’s, Tukey’s, or Dunnett’s multiple comparisons test or two-way ANOVA followed by Tukey’s multiple comparisons test. *P* value less than 0.05 denoted the presence of a statistically significant difference. All bars in dot plot figures represent means.

## Results

### Loss of Gcn2 did not induce either spontaneous PVOD or PH in mice

To determine if the KO mice develop spontaneous PVOD or PH, we carried out hemodynamic measurements and histological assessment of pulmonary-veno occlusion. As shown in **Figure 1A**, *Gcn2* mRNA expression was abolished in KO lungs, but the expression of PERK, another stress response kinase^28,36^ was not affected, demonstrating gene-specific knockout. At ages of 3.5, 12 and 23 mos., right ventricle systolic pressure (RVSP) of KO mice was 23.7±1.3, 23.5±1.3, 24.2±0.8, respectively, compared to 22.6±1.9 of 3.5 mos. old WT mice (**Figure IA in the Data Supplement**). The RV/Left Ventricule+Septum (RV/LV+S) weight ratio, an indicator of RV hypertrophy, of KO mice was also similar to WT mice (**Figure IB in the Data Supplement**). These data demonstrate that KO mice do not develop spontaneous PH. Furthermore, H&E staining and Russell-Movat pentachrome staining of these mouse lung sections revealed no evidence of vascular occlusion or luminal narrowing (**Figure II in the Data Supplement**), illustrating that KO mice do not have the histopathological features of PVOD.

**Figure 1.**
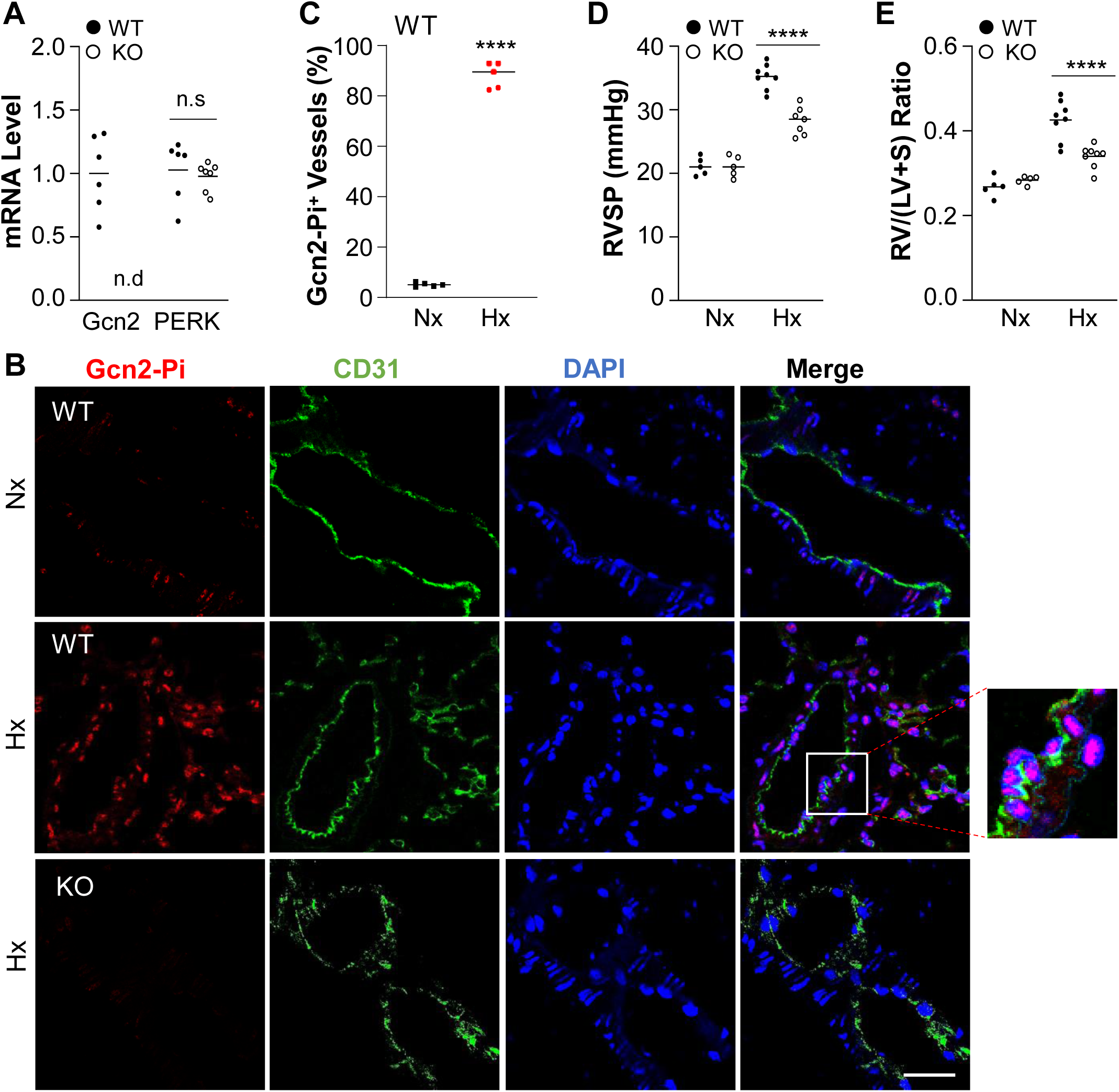
GCN2 deficiency attenuated hypoxia-induced PH in mice. (**A**) Quantitative RT-PCR analysis confirming GCN2-specific disruption in *Eif2ak4^-/-^* (KO) mouse lungs. n.d, not detected. (**B**) Representative micrographs of anti-phopho-GCN2 immunostaining of mouse lungs demonstrating prominent Thr898 phosphorylation-GCN2 (GCN2-Pi) in pulmonary vascular ECs in hypoxic WT but not KO mice. Lung tissue cryosections from normoxic and hypoxic (3 weeks) WT mice and hypoxic KO mice were immunostained with anti-Thr phospho-GCN2 antibody (red). ECs were immunostained with anti-CD31 (green) and nuclei were counterstained with DAPI (blue). Scale bar, 50µm. (**C**) Quantification of GCN2-Pi-positive vessels. Data are expressed as percentage of positive vessels in each mouse lung. (**D**) RVSP measurement showing a marked increase of RVSP in hypoxic WT mice compared to normoxic WT mice, which was markedly reduced in hypoxic KO mice. (**E**) RV hypertrophy evident by increased RV/LV+S ratio seen in hypoxic WT mice was markedly reduced in hypoxic KO mice. Nx=normoxia, Hx=hypoxia. Bars represent means. n=4female (F)+4male (M) hypoxic WT, 4F+4M hypoxic KO, 3F+2M normoxic WT and 3F+2M normoxic KO mice were used in (**D**, **E**). n.s., not significant. ****, P < 0.0001. Unpaired two-tailed t test (**C**); Two-way ANOVA with Tukey’s multiple comparisons test (**D**, **E**).

### Loss of Gcn2 protected mice from hypoxia-induced PH

Since KO mice do not spontaneously develop PVOD or PH, we challenged the mice with chronic hypoxia to assess their responses to PH development. Immunofluorescent staining and Western blotting revealed prominent Gcn2 phosphorylation at Thr898 indicative of activation^22^ in pulmonary vascular endothelial cells (ECs) of hypoxic WT mice, which was completely absent in KO lungs (**Figure 1B** and **1C** and **Figure III in the Data Supplement**). Following 3 weeks of hypoxia (10% O_2_), WT mice developed PH evident by marked increase of RVSP (35.2±1.9 mmHg), whereas KO mice exhibited a marked lower RVSP (28.2±2.2 mmHg) (**Figure 1D**). Hypoxic KO mice also had decreased RV/LV+S ratio, i.e. RV hypertrophy, compared to hypoxic WT mice (**Figure 1E**). These data demonstrate that loss of Gcn2 kinase function inhibits hypoxia-induced PH in mice.

Next, we performed pulmonary pathology examination by Russell-Movat pentachrome staining and anti-α-smooth muscle actin (α-SMA) immunofluorescent staining of these lung tissues. Russell-Movat pentachrome staining showed that hypoxia-induced increase in pulmonary vessel media wall thickness in WT mice was attenuated in KO mice (**Figure 2A through 2C**). The number of muscularized distal pulmonary arterioles shown as α-SMA-positive staining was also markedly reduced in hypoxic KO mice compared to hypoxic WT mice (**Figure 2D** and **2E**). To determine whether Gcn2 mediates pulmonary vascular EC and SMC proliferation, we assessed cell proliferation by anti-Ki67 immunostaining. After one-week hypoxia exposure, WT mice showed a 6-fold increase in the number of proliferating SMCs (i.e., Ki67^+^/α-SMA^+^ cells) (**Figure 2F** and **2G**) and a 15-fold increase of Ki67^+^/CD31^+^ cells (proliferating ECs) (**Figure IV in the Data Supplement**), respectively, compared to those under normoxia condition. However, hypoxic KO mice exhibited a marked reduction in the number of proliferating SMCs and ECs compared to hypoxic WT mice (**Figure 2F** and **2G**, and **Figure IV in the Data Supplement**). Together, these data suggest that GCN2 activation by hypoxia contributes to pulmonary vascular remodeling in hypoxic WT mice. Together, these data suggest that GCN2 activation by hypoxia contributes to pulmonary vascular remodeling in hypoxic WT mice.

**Figure 2.**
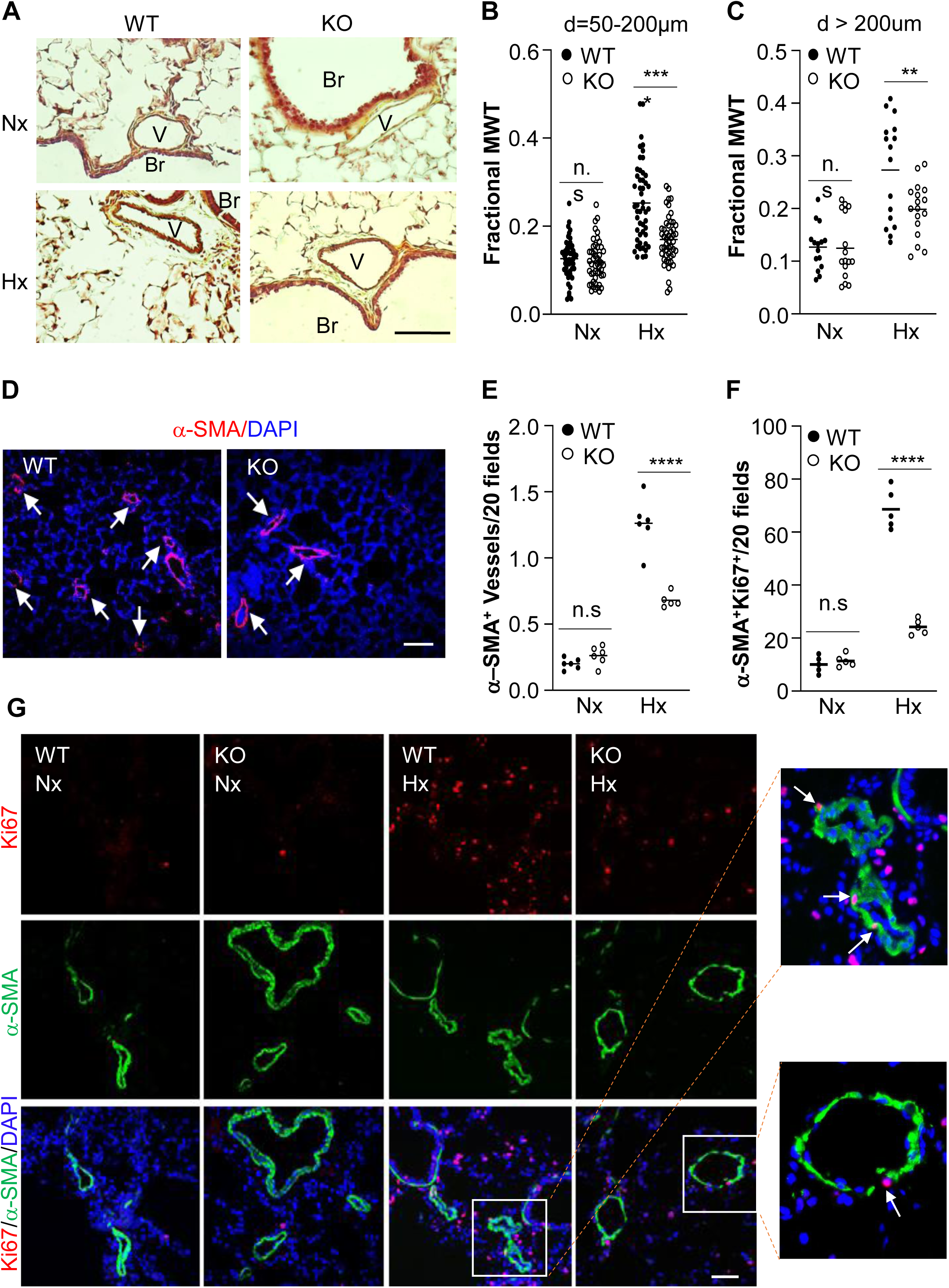
Reduced pulmonary vascular remodeling in hypoxic KO mice compared to hypoxic WT mice. (**A**) Representative micrographs of Russell-Movat pentachrome staining of mouse lung sections. Br, bronchiole; V, vessel. Scale bar, 50 µm. (**B, C**) Quantification of pulmonary vessel wall thickness of different diameter (d) sizes of vessels. 50 vessels/group and 5 mice/group for (**B**); 15-16 vessels/group and 5 mice/group for (**C**). MWT=media wall thickness. (**D**) Representative micrographs of anti-α-SMA staining of hypoxic WT and hypoxic KO lung sections showing reduced number of muscularized distal pulmonary vessels in hypoxic KO lungs. Nuclei were counterstained with DAPI (blue). Arrows point to muscularized vessels. Scale bar, 50 µm. (**E**) Quantification of muscularized distal pulmonary vessels. The total number of α-SMA-positive distal pulmonary vessels (d ≤ 50µm) of 20× fields of each section was used for each mouse. 20 fields of each mouse section, 5 mice for each group. (**F**, **G**) Quantification of SMC proliferation in mouse lungs. Representative micrographs of immunostaining of mouse lung sections were shown in (**G**). Mouse lungs were collected from 3.5 months old mice under normoxia or 1 week hypoxia for sectioning and immunostaining with anti-Ki67 (red) to identify proliferative cells. SMCs were immunostained with anti-α-SMA (green). Nuclei were counterstained with DAPI (blue). Scale bar, 50 µm. Bars represent means. n.s=not significant. **, P < 0.01; ****, P < 0.0001. Two-way ANOVA with Tukey’s multiple comparisons test (**B**, **C**, **E, F**).

### GCN2 is phosphorylated and activated via PDK1 in response to hypoxia

To determine if GCN2 is activated in response to hypoxia *in vitro*, we challenged primary culture of HLMVECs with hypoxia (1% O_2_). GCN2 was highly activated, evident by immunofluorescent staining of Thr899 hyperphosphorylation in hypoxic HLMVECs compared to normoxic cells (**Figure 3A**). It was also observed that GCN2 was activated in the cytosol in response to hypoxia within 2 hours exposure and continued to be activated but concentrated in the nucleus at 4 and 8 hours hypoxia exposure. Western blotting also confirmed the increase of GCN2 activity by hypoxia, evident by Thr899 phosphorylation in GCN2 and phosphorylating its downstream target EIF2a at Ser51 site (**Figure 3B through 3F**). Given that Gcn2 can be phosphorylated by Snf1 [the ortholog of mammalian AMP-activated kinase (AMPK)] under histidine starvation condition in yeast^37^ or yeast Pkh1 (the ortholog of mammalian PDK1) *in vitro*, ^38^ we next determined whether hypoxia-activated GCN2 was mediated by either pathway. Hypoxic HLMVECs were treated with either AMPK inhibitor compound C or PDK1 inhibitor GSK233470.^39^ Western blotting demonstrated that hypoxia induced GCN2 phosphorylation through PDK1, evident by decreased GCN2 phosphorylation level by PDK1 inhibitor GSK 2334470 in a dose-dependent manner (**Figure 3G through 3I**) but not by AMPK inhibition (**Figure V in the Data Supplement**). In contrast, compound C treatment further increased GCN2 phosphorylation.

**Figure 3.**
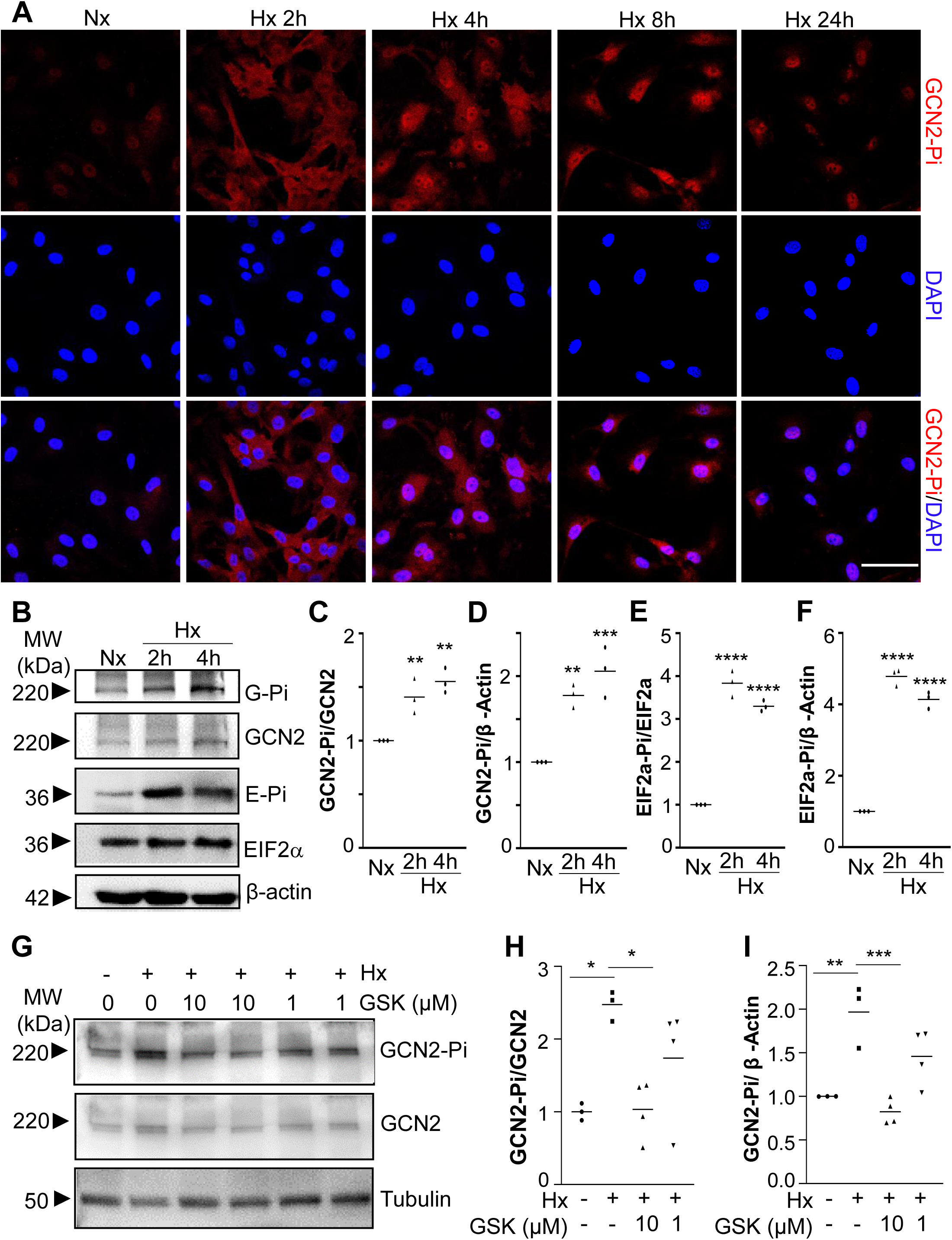
Hypoxia induces GCN2 phosphorylation and activation. (**A**) Representative micrographs of anti-phospho-GCN2 immunostaining showing GCN2 hyperphosphorylation by hypoxia challenge in primary cultures of HLMVECs. Fixed cells at indicated times post-hypoxia (1% O_2_) challenge (Hx) or normoxia (Nx) were immunostained with anti-Thr899phospho-GCN2 (red). Nuclei were counterstained with DAPI (blue). (**B-F**) Western blotting demonstrating hypoxia induced GCN2 phosphorylation and activation. HLMVECs were lysed for Western blotting with anti-Thr899 Phospho-GCN2 (GCN2-Pi) antibody and anti-Ser51 Phospho-EIF2a (E-Pi) antibody. Total GCN2 and EIF2a levels were assessed by anti-GCN2 antibody and anti-EIF2a antibody while anti-β-actin was used as a loading control (**B**). The band intensities of phosphor-GCN2 and phosphor-EIF2a (EIF2a-Pi) were quantified **(C-F)**. (**G-I**) Western blotting demonstrating hypoxia-induced GCN2 phosphorylation was inhibited by PDK1 inhibitor (GSK=GSK2334470) treatment in a dose-dependent manner. HLMVECs in complete growth medium were treated with GSK2334470 at 10 or 1 µM or control vehicle under 2h hypoxia challenge (1% O_2_), then cells were lysed for Western blotting with anti-Thr899 Phospho-GCN2 (GCN2-Pi) antibody. Total GCN2 levels were assessed by anti-GCN2 antibody while anti-β-actin was used as a loading control (**G**). The band intensities of phosphor-GCN2 were quantified **(H, I)**. *, P < 0.05; **, P < 0.01; ***, P < 0.001; ****, P < 0.0001. One-way ANOVA with Dunnett’s multiple comparisons test (**C-F**), with Sidak’s multiple comparisons test (**H**), and with Tukey’s multiple comparisons test (**I**).

### Hypoxia-induced Endothelin-1 (Edn1) expression is suppressed by GCN2 deficiency

To determine the molecular mechanism of GCN2 in regulating PH and vascular remodeling, we performed whole-transcriptome RNA sequencing analysis of both WT and KO mouse lung tissues subjected to normoxia and hypoxia conditions (**Figure 4A through 4C**). RNA sequencing data analysis showed that upregulated genes in WT lungs with hypoxia challenge such as *Angptl4*, *Edn1*, *CD59a*, *Tusc5*, *Fkbp5* and *Pde4d* were completely normalized in KO lungs with hypoxia challenge, and downregulated genes in WT lungs with hypoxia challenge such as *Lars2*, *Exosc5*, *Muc5b*, and some ribosome protein genes (*Rrp*s) were completely normalized in KO lungs with hypoxia challenge. Endothelin-1 (Edn1), a potent vasoconstrictor in the pathogenesis of PAH,^40–43^ is ranked at the top 2 gene of the upregulated gene list, suggesting Edn1 as the potential downstream target gene of GCN2 in regulating PH. Quantitative RT-PCR analysis and Western blotting confirmed marked increases in the expression of Edn1 in WT mouse lungs with hypoxia compared to those with normoxia, whereas markedly reduced in KO mouse lungs with hypoxia (**Figure 4D** and **4E**).

**Figure 4.**
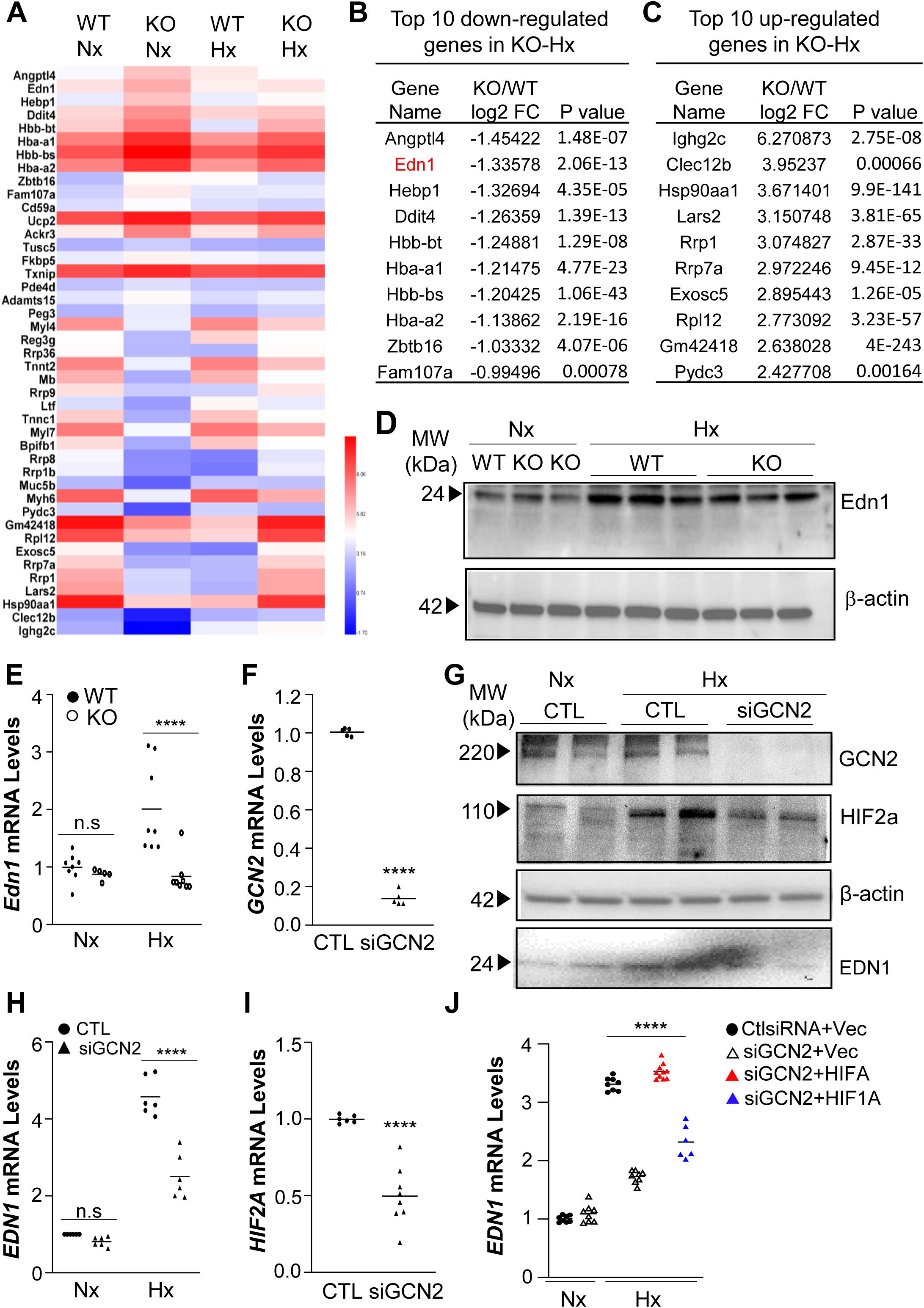
Hypoxia induces Edn1 expression through GCN2 both in mouse lungs and human lung ECs. (**A**) Representative heatmap of RNA sequencing analysis of WT normoxia (WT Nx), WT hypoxia (WT Hx), KO normoxia (KO Nx), and KO hypoxia (KO Hx) mouse lungs (n=4 mice combined per group). (**B**) List of top ten upregulated genes by hypoxia in WT mouse lungs that were significantly reduced in KO hypoxia mouse lungs. (**C**) List of top ten downregulated genes by hypoxia in WT mouse lungs that were significantly increased in KO hypoxia mouse lungs. (**D**) Western blotting demonstrating Edn1 protein levels were upregulated in lung tissues of hypoxic WT mice while reduced in hypoxic KO mice. (**E**) Quantitative RT-PCR analysis confirming *Edn1* mRNA upregulation in WT hypoxia mouse lungs while reduced in KO hypoxia mouse lungs. (**F**) Quantitative RT-PCR analysis demonstrating GCN2 siRNA (siGCN2)-mediated knockdown of GCN2 in primary culture of HLMVECs. CTL, control RNA oligonucleotides. (**G**) Western blotting confirmation of reduced HIF-2α and EDN1 expression in GCN2-deficient HLMVECs compared to control cells under hypoxia exposure. (**H**) Quantitative RT-PCR analysis demonstrating hypoxia-induced *EDN1* mRNA expression was mediated by GCN2 in HLMVECs. (**I**) Quantitative RT-PCR analysis demonstrating inhibited *HIF2A* mRNA expression in GCN2-deficient HLMVECs. (**J**) Quantitative RT-PCR analysis demonstrating GCN2 mediated *EDN1* expression through HIF-2α rather than HIF-1α. HLMVECs were transfected with either GCN2 siRNA (siGCN2) or control (CTL) and *HIF1A* or *HIF2A* plasmid combination or vector plasmid as control and then challenged with hypoxia or remained in normoxia. At 48 h post-hypoxia challenge (Hx) or normoxia (Nx), the cells were lysed for RNA and protein isolation for quantitative RT-PCR and Western blotting analysis, respectively. n.s., not significant. ****, P < 0.0001. Unpaired two-tailed t test (**F**); Two-way ANOVA with Tukey’s multiple comparisons test (**E, H-J**).

To further determine the effect of GCN2 deficiency on Edn1 expression, which is predominantly expressed in ECs, we knocked down GCN2 with *GCN2* siRNA in HLMVECs and exposed the cells to hypoxia. Quantitative RT-PCR analysis demonstrated a 90% knockdown efficiency of GCN2 (**Figure 4F**). As expected, hypoxia (1% O_2_) exposure induced a marked increase of EDN1 protein expression and ∼4-fold increase in *EDN1* mRNA expression in control cells, while they were markedly reduced in GCN2-deficient cells, demonstrating that hypoxia induces EDN1 expression through GCN2 (**Figure 4G** and **4H**). Since EDN1 is a downstream target of hypoxia-inducible factor (HIF) 1,^44,45^ we further determined whether GCN2 regulates EDN1 through HIFs. Quantitative RT-PCR analysis showed inhibited mRNA levels of *HIF2A* in hypoxic GCN2-deficient HLMVECs compared to control cells (**Figure 4H**). Western blotting also demonstrated that hypoxia-induced increases of HIF-2α protein levels were inhibited in GCN2-deficient HLMVECs (**Figure 4G**). However, hypoxia-induced increases of HIF-1α protein levels were largely not reduced in GCN2-deficient HLMVECs (**Figure VIA in the Data Supplement**). To determine if HIF-1α or HIF-2α overexpression can rescue EDN1 expression in GCN2-deficient HLMVECs, GCN2 siRNA or control siRNA and either *HIF2A* plasmid, *HIF1A* plasmid or vector plasmid were co-transfected to HLMVECs, and the cells were exposed to hypoxia. Quantitative RT-PCR analysis showed that hypoxia-induced *EDN1* expression was inhibited by GCN2 deficiency, which was fully rescued by *HIF2A* plasmid-mediated overexpression of HIF-2α. However, HIF-1α overexpression only partially rescued EDN1 expression (**Figure 4J** and **Figure VIB and VIC in the Data Supplement**), demonstrating that GCN2 regulates EDN1 expression through regulating HIF-2α expression rather than HIF-1α.

### Restored endothelial Edn1 expression in KO mice reversed the reduced PH phenotype

We next performed rescue study *in vivo* employing the endothelium-targeted nanoparticle delivery of gene^35^ to determine if restoration of endothelial Edn1 expression in Gcn2-deficient mice can reverse the reduced phenotype of PH in response to hypoxia. Mixture of EndoNP1 nanoparticles:plasmid DNA expressing mouse *Edn1* under the control of human *CDH5* promoter, or *GFP* plasmid (as a control) was administered retro-orbitally into KO mice weekly. WT mice also received *GFP* plasmid (**Figure 5A**). After three weeks exposure to hypoxia (10% O_2_), mouse lung ECs were isolated for Western blotting. Edn1 was highly expressed in KO mice with *Edn1* plasmid administration compared to those with *GFP* control plasmid DNA (**Figure 5B**). Hemodynamic measurements demonstrated that reduced RVSP and RV/(LV+S) ratio seen in KO mice with GFP plasmid DNA were reversed in *Edn1* plasmid DNA-administered KO mice (**Figure 5C** and **5D**). α-SMA immunostaining and histological analysis also showed that reduced number of muscularized distal pulmonary arterioles and decreased pulmonary arterial medium wall thickness in KO mice with *GFP* plasmid DNA were also rescued in *Edn1* plasmid DNA-administered KO mice (**Figure 5E through 5H**). Taken together, these data demonstrated that decreased endothelial Edn1 expression in KO mice plays a causal role in inhibiting hypoxia-induced PH.

**Figure 5.**
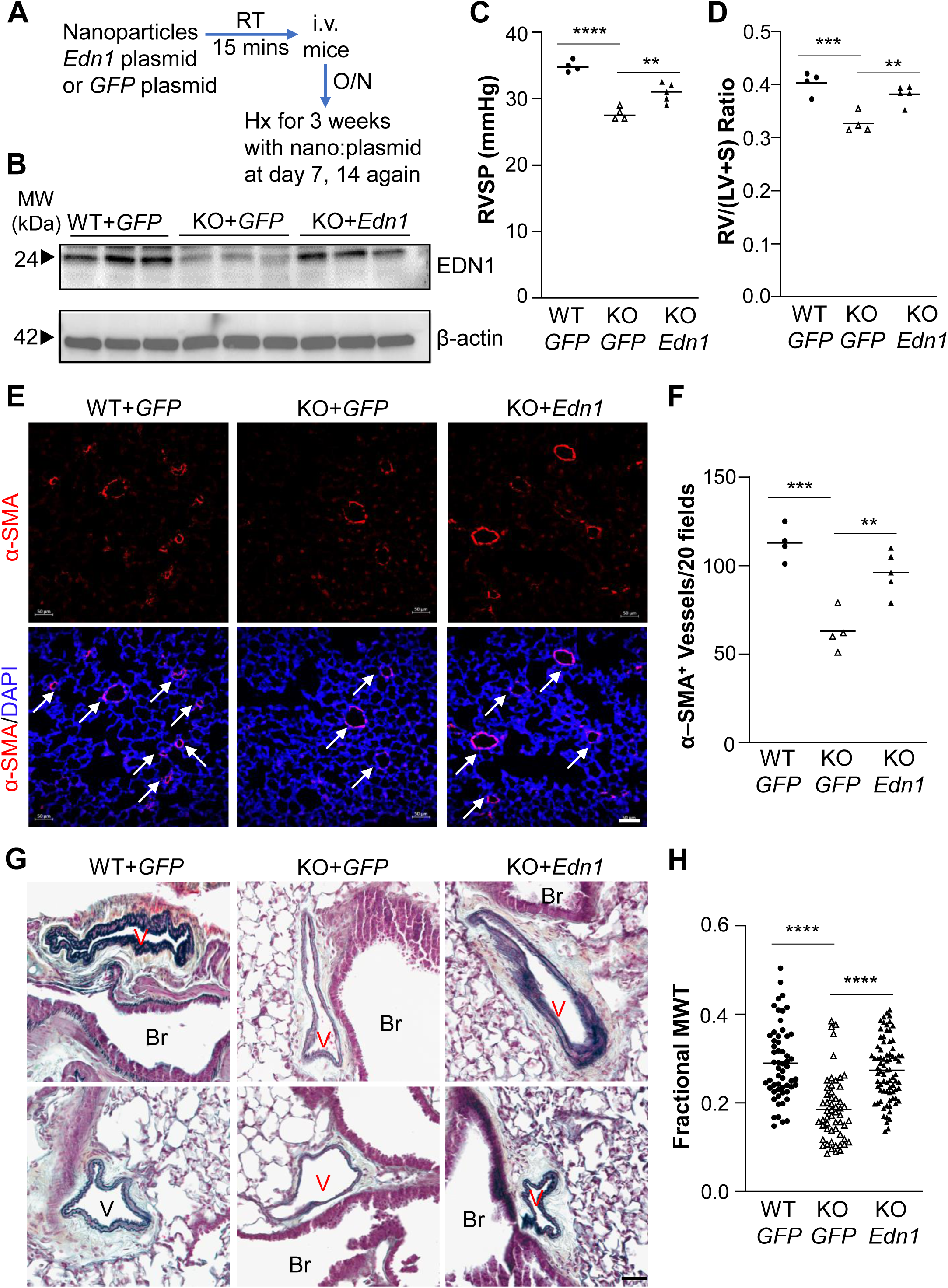
Restored expression of endothelial Edn1 in GCN2-deficient mice reversed the reduced PH phenotype. (**A**) Diagram showing the experimental procedure to overexpress endothelial *Edn1* in *Gcn2*-deficient mice. Mixture of nanoparticles:plasmid DNA expressing *Edn1* or *GFP* (control) under the control of *CDH5* promoter was administered retro-orbitally to 11 weeks old KO mice. WT mice received *GFP* plasmid. After overnight (16h), the mice were subjected to hypoxia. Nanoparticles:plasmid DNA mixtures were administered weekly for total 3 doses. (**B**) Western blotting demonstrating restored Edn1 expression in KO mice with *Edn1* plasmid administration compared to WT and KO mice with *GFP* plasmid. (**C**) RVSP measurement showing reduced PH in KO mice with *GFP* plasmid was reversed in KO mice with *Edn1* plasmid. (**D**) RV/(LV+S) ratio also showing reduced RV hypertrophy in KO mice with *GFP* plasmid was markedly rescued with restored *Edn1* expression. (**E**) Representative micrographs of anti-α-SMA staining of lung sections showing reduced muscularization of distal pulmonary vessels in hypoxic KO+*GFP* lungs was reversed in hypoxic KO+*Edn1* lungs. Nuclei were counterstained with DAPI (blue). Arrows point to muscularized vessels. (**F**) Quantification of muscularized distal pulmonary vessels. (**G**) Representative micrographs of Russell-Movat pentachrome staining of mouse lung sections. Br, bronchiole; V, vessel. (**H**) Quantification of pulmonary vessel media wall thickness. n=2M+2F WT+*GFP*, 2M+2F KO+*GFP*, and 2M+3F KO+*Edn1*. Scale bars, 50 µm. **, P < 0.01; ***, P < 0.001; ****, P < 0.0001. One-way ANOVA with Tukey’s multiple comparisons test (**C, D, F, H**).

### Loss of Gcn2 in ECs in mice attenuates hypoxia-induced PH

We next generated EC-specific knockout mice through nanoparticle delivery of the CRISPR/Cas9 system targeting the GCN2 kinase domain to determine if loss of endothelial Gcn2 in mice has a similar phenotype (**Figure 6A**). Mixture of EndoNP1 nanoparticles:CRISPR plasmid DNA expressing Cas9 under the control of *CDH5* promoter and *Eif2ak4*-specific guide RNA (gRNA) driven by *U6* promoter were administered to C57BL/6 WT mice at age of 10 weeks and plasmid expressing control gRNA was administered to a separate cohort of WT mice as WT controls. Immunofluorescent staining of GCN2-Pi demonstrated loss of Gcn2 selectively in vascular ECs in *Eif2ak4* gRNA plasmid-administered mice under hypoxia exposure (**Figure 6B**). Hemodynamic measurements showed that loss of endothelial Gcn2 resulted in reduced PH phenotype, evident by decreased RVSP and RV/(LV+S) ratio, compared with control plasmid-administered mice (**Figure 6C** and **6D**). Russell-Movat pentachrome staining and immunofluorescent staining of α-SMA also demonstrated that loss of endothelial Gcn2 in mice attenuated pulmonary vascular media wall thickness and reduced the number of muscularized distal pulmonary vessels (**Figure 6E through 6H**). Therefore, loss of endothelial Gcn2 in mice attenuated hypoxia-induced PH.

**Figure 6.**
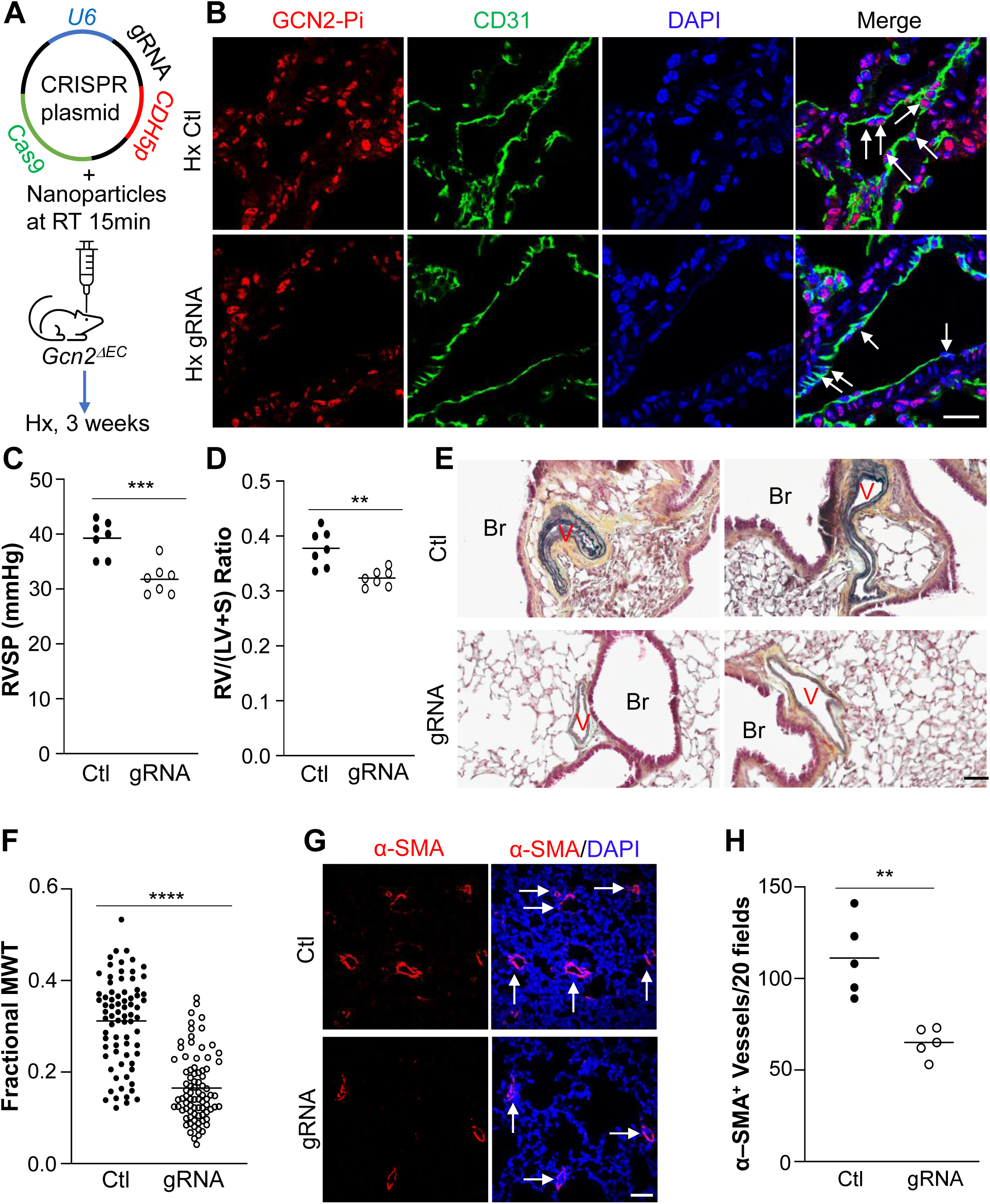
Loss of endothelial GCN2 in mice attenuates hypoxia-induced PH. (**A**) Diagram showing the strategy to generate EC-specific *Gcn2* knockout (*Gcn2^ΔEC^*) mice through nanoparticles-mediated delivery of CRISPR/*Eif2ak42* guide RNA (gRNA). C57BL/6 WT mice were administered retro-orbitally with mixture of nanoparticles:CRISPR plasmid DNA expressing Cas9 under the control of *CDH5* promoter and *Eif2ak4* gRNA driven by *U6* promoter or control plasmid expressing control gRNA at 10 weeks of age and then exposed to hypoxia at 11 weeks of age for 3 weeks. (**B**) Representative micrographs of anti-phopho-GCN2 immunostaining of mouse lungs showing prominent Thr898 phosphorylation-GCN2 (GCN2-Pi) (red) in pulmonary vascular ECs in hypoxic WT mice while diminished in *Gcn2* gRNA-administered mice, demonstrating loss of GCN2 selectively in vascular ECs. Lung tissue cryosections were counterstained with anti-CD31 antibody (green) and DAPI (blue). Arrows point to EC nuclei. Scale bar, 20 µm. (**C, D**) RVSP and RV/(LV+S) ratio measurements showing reduced PH and RV hypertrophy in *Gcn2* gRNA-administered mice compared to control gRNA mice. (**E, F**) Representative micrographs of Russell-Movat pentachrome staining of hypoxic control mouse lung sections and hypoxic *Gcn2* gRNA mouse lung sections and quantification of pulmonary media wall thickness. Br, bronchiole; V, vessel. Scale bar, 50 µm. (**G, H**) Representative micrographs of anti-α-SMA immunostaining of hypoxic control mouse lung sections and hypoxic *Gcn2* gRNA mouse lung sections and quantification of distal pulmonary arterial muscularization. n=3F+4M control, and 4F+3M *Gcn2* gRNA. Arrows point to muscularized vessels. Scale bar, 50 µm. **, P < 0.01; ***, P < 0.001; ****, P < 0.0001. Unpaired two-tailed t test (**C, D, F, H**).

### Pharmacological inhibition of GCN2 suppresses PAH in monocrotaline rats

To determine if GCN2 is a potential therapeutic target for PAH treatment, we first did immunofluorescent staining with anti-Thr899 phosphor-GCN2 in rat lungs challenged with monocrotaline (MCT). GCN2 was hyperphosphorylated in vascular ECs in MCT rat lungs in contrast to naïve control rats, suggesting GCN2 hyperactivation in response to MCT challenge. We next determined if pharmacological inhibition of GCN2 could inhibit PAH in MCT rats. At 2 weeks after MCT challenge, the rats were treated with either GCN2 inhibitor A-92 ^46^ (**Figure VIIA in the Data Supplement**) or vehicle. Western blotting demonstrated MCT-induced GCN2 activation in rat lung tissue was inhibited by A-92 treatment (**Figure VIIB in the Data Supplement**). Accordingly, MCT-induced increases in RVSP and RV/(LV+S) ratio were markedly reduced in A-92-treated rats compared to vehicle-treated ones (**Figure 7C** and **7D**, and **Figure VIIC** and **VIID in the Data Supplement**). A-92 treatment also attenuated pulmonary vascular remodeling including media wall thickening and the number of muscularized distal pulmonary vessels in MCT rats (**Figure 7E through 7H**). A-92 treatment also reduced vascular SMC and EC proliferation in MCT rats (**Figure ⅤIII in the Data Supplement**). Together, these data suggest that Inhibition of GCN2 activation suppresses pulmonary vascular remodeling and PH supporting GCN2 as a potential therapeutic target for PAH treatment.

**Figure 7.**
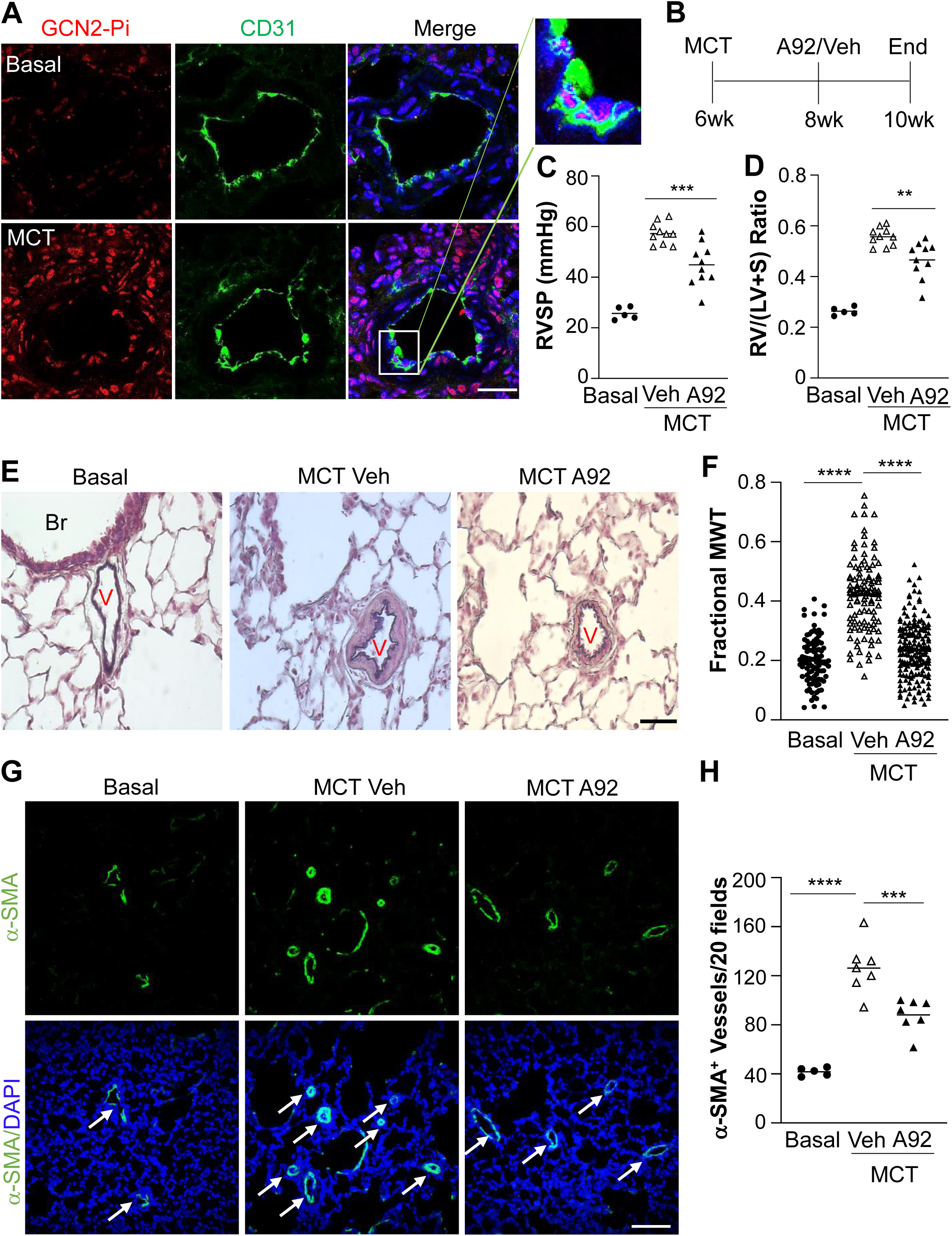
GCN2 inhibition attenuates MCT-induced PAH in rats. (A) Representative micrographs of anti-phopho-GCN2 immunostaining of rat lungs demonstrating prominent Thr898 phosphorylation-GCN2 (GCN2-Pi) in pulmonary vascular ECs in MCT rats. Lung tissue cryosections from basal control rats and MCT-treated rats (4 weeks) were immunostained with anti-Thr898 phospho-GCN2 antibody (red). ECs were immunostained with anti-CD31 (green) and nuclei were counterstained with DAPI (blue). Scale bar, 50µm. (**B**) Graphic presentation of the experimental procedure. A-92 (0.5 mg/kg, i.p. daily) was administered to rats at 2 weeks post-MCT. A92=A-92; Veh=vehicle. (**C**)RVSP measurement showing a marked increase of RVSP in MCT rats treated with vehicle compared to basal rats, which was markedly reduced in A-92-treated MCT rats. (**D**) RV hypertrophy evident by increased RV/LV+S ratio seen in MCT vehicle rats was markedly reduced in A-92-treated MCT rats. (**E**) Representative micrographs of Russell-Movat pentachrome staining of rat lung sections showing attenuated vessel wall thickening. V, vessel. (**F**) Quantification of pulmonary vessel wall thickness. MWT=media wall thickness. n=96 vessels with 5 control rats, n=107 vessels with 5 vehicle-treated MCT rats, n=174 vessels with 5 A-92-treated rats. (**G**) Representative micrographs of anti-α-SMA (green) staining of basal rat, MCT vehicle rat and A-92-treated MCT rat lung sections showing reduced muscularization of distal pulmonary vessels in A-92-treated MCT rat lungs. Nuclei were counterstained with DAPI (blue). Arrows point to muscularized distal pulmonary vessels. (**H**) Quantification of muscularized distal pulmonary vessels. The total number of α-SMA-positive distal pulmonary vessels (d ≤ 50µm) of 20× fields of each section was used for each rat. Arrows point to muscularized vessels. Bars represent means. n=5M basal, 10M MCT rats+vehicle, and 10M MCT rats+A-92. Scale bars, 50µm. **, P < 0.01; ***, P < 0.001; ****, P < 0.0001. One-way ANOVA with Dunnett’s multiple comparisons test (**C, D, F, H**).

### GCN2 was highly activated in pulmonary vascular ECs of IPAH patients without marked changes of protein expression

To address the clinical relevance of GCN2 activation in the pathogenesis of PAH, we first quantified GCN2 mRNA expression in lung tissues from healthy donors and IPAH patients (**Table Ⅰ in the Data Supplement**) by quantitative RT-PCR analysis. Although GCN2 mRNA expression varied among individuals in both control donor group and IPAH patient group, there was no significant difference in the mRNA levels between the 2 groups (**Figure IX in the Data Supplement**). Western blotting also revealed similar GCN2 protein levels in lungs of donors and PAH patients (**Figure 8A** and **8B**). Similarly, there was no significant difference in GCN2 protein levels in pulmonary arterial ECs of donors and IPAH patients (**Figure 8C** and **8D**). We next determined whether GCN2 phosphorylation and activation differs between donors and patients. Western blotting analysis showed significantly increased phosphorylation of GCN2 in whole lung tissues of IPAH patients compared to control donors (**Figure 8E**). Lung sections were immunostained with anti-Thr899 Phospho-GCN2 and anti-VWF to assess GCN2 activation in pulmonary vascular ECs. The number of GCN2 phosphorylated vascular ECs in PAH patient lungs was ∼ 5-fold higher than the number of those in donor lungs, suggesting highly activated GCN2 in pulmonary vasculature of IPAH patients (**Figure 8F** and **8G**). These data support the concept that GCN2 activation in ECs contributes to pulmonary vascular remodeling and PAH development in patients.

**Figure 8.**
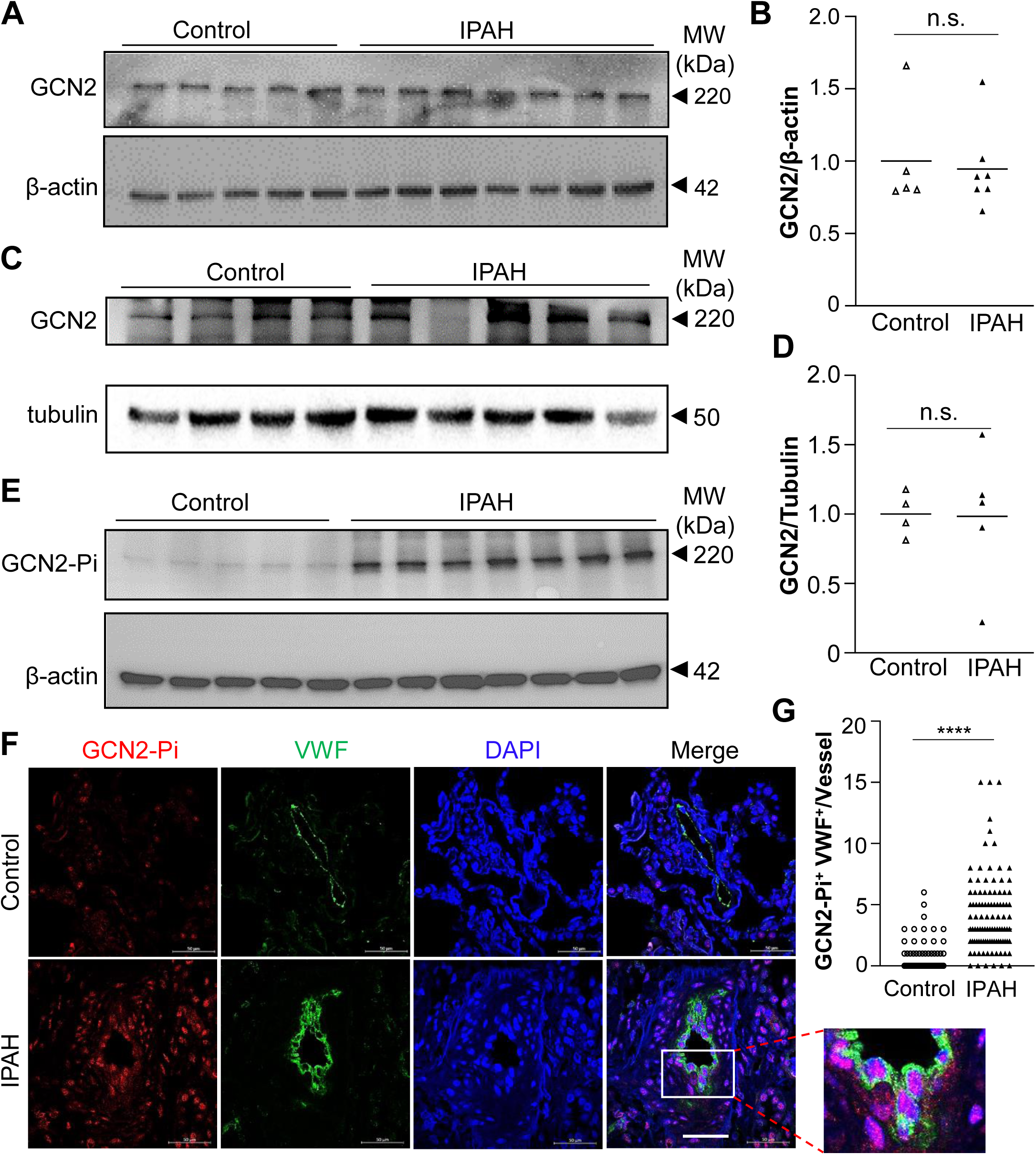
Prominent GCN2 phosphorylation/activation in ECs of pulmonary vascular lesions of IPAH patients. (**A**, **B**) Western blotting of anti-GCN2 and quantification demonstrating no difference in GCN2 total protein expression in whole lung tissues of IPAH patients compared to normal donors. **(C, D)** Western blotting of anti-GCN2 and quantification demonstrating no difference in GCN2 total protein expression in pulmonary arterial endothelial cells of IPAH patients compared to normal donors. **(E)** Western blotting demonstrating extensive GCN2 phosphorylation in IPAH patient lung tissues but minimal in control donor lung tissues. (**F, G**) Prominent GCN2 phosphorylation in ECs of pulmonary vascular lesions of IPAH patients but not in normal donor lungs. Formalin-fixed lung sections from 5 IPAH patients and 5 non-PAH donors were immunostained with anti-Thr899phospho-GCN2 (GCN2-Pi) (red) and anti-vWF (green). Representative micrographs of immunofluorescent staining were shown in (**F**). The number of GCN2-Pi^+^ ECs in each vessel was quantified (15 vessels/sample) (**G**). Scale bar, 50µm. n.s., not significant. ****, P < 0.0001. Mann-Whitney U test (**B, G**). Unpaired two-tailed t-test (**D**).

## Discussion

The present study has defined the role of GCN2 activation in mediating pulmonary vascular remodeling and PH development. We observed no spontaneous PVOD and PH in KO mice. However, loss of GCN2 inhibited hypoxia-induced pulmonary vascular remodeling and PH in mice. Mechanistic studies demonstrated hypoxia activated GCN2 via PDK1 in HLMVECs, which mediated hypoxia-induced EDN1 expression via HIF-2α. Restored Edn1 expression in mouse lung ECs rescued the reduced PH phenotype of KO mice in response to hypoxia. Additionally, knockout of endothelial GCN2 in mice attenuated hypoxia-induced PH. Moreover, inhibition of GCN2 activity through GCN2 inhibitor A-92 treatment inhibited MCT-induced PAH and pulmonary vascular remodeling in rats. In IPAH lungs, GCN2 phosphorylation was markedly increased in pulmonary vascular ECs without marked changes of GCN2 expression. Together, these data demonstrate the requisite role of GCN2 activation in mediating pulmonary vascular remodeling and PAH development. Thus, targeting GCN2 signaling is a novel therapeutic strategy for treatment of PAH without GCN2 loss-of-function mutations.

Published studies have demonstrated a strong relationship of loss-of-function mutations of *EIF2AK4* and PVOD development in patients. To our surprise, loss of GCN2 in mice doesn’t induce spontaneous PVOD. Given that the KO mice used in this study have genetic deletion of exon 12 of the *Eif2ak4* gene encoding amino acids 606-648, an essential region for its kinase activity,^14^ it is possible that the N-terminal 605 amino acids may retain some of the GCN2 function domains which are sufficient to inhibit PVOD development. Among 41 reported variants of *EIF2AK4* gene associated with classic PAH (i.e., non-PVOD PAH), none of them are located in exon 12 of the gene;^47^ while among 32 reported variants of *EIF2AK4* associated with PVOD and/or pulmonary capillary haemangiomatosis (PCH), only 2 of them are located in exon 12, including 1 missense variant at amino acid 643,^29^ suggesting that most of the reported mutations of *EIF2AK4* associated with PAH or PVOD are not related to exon 12 in the essential kinase domain. This may also explain our observation that deletion of exon 12 in GCN2 kinase domain does not induce spontaneous PAH or PVOD. Comparing all the reported variants of *EIF2AK4* between classic PAH and PVOD patients, only 3 variants are shared,^47^ indicating that mutations in different functional domains of GCN2 may play distinct roles in the development of classic PAH and PVOD. It is possible that a second “hit” coupled with loss of GCN2 is required for PVOD development.

Our studies show GCN2 hyperphosphorylation of Thr898 at the kinase domain which indicates GCN2 activation^21–25^ in pulmonary vascular ECs of hypoxic WT mice. We also observed rapid and extensive phosphorylation of GCN2 in cultured HLMVECs in response to hypoxia and GCN2 is required for hypoxia-induced vascular cell proliferation in mice. Thus, hypoxia-induced activation of GCN2 plays a causal role in mediating pulmonary vascular remodeling and PH development. Surprisingly, we also observed an extensive GCN2 phosphorylation in pulmonary vascular ECs in IPAH patients, demonstrating the clinical relevance of GCN2 activation in the pathogenesis of PAH. There are contradictory reports about GCN2 protein levels in IPAH lungs. One report shows diminished GCN2 protein levels in IPAH lungs and monocrotaline-treated rat lungs, indicating loss of GCN2 induces PAH.^48^ Another report, however, demonstrates similar GCN2 protein levels in IPAH and control lungs.^49^ Our study here shows no marked difference in GCN2 expression, both mRNA and protein levels, in whole lung tissues as well as pulmonary vascular ECs between IPAH patient lungs and control lungs. Furthermore, our study provides unequivocal evidence of extensive GCN2 phosphorylation in pulmonary vascular lesions of IPAH patients, especially in vascular ECs.

These data demonstrate that GCN2 activation, rather than GCN2 expression changes, plays a crucial role in pulmonary vascular remodeling and PAH development in patients. In response to amino acid starvation, GCN2 is activated by the binding of uncharged tRNAs accumulated in amino acid-starved cells to the histidyl-tRNA synthetase (HisRS)-related domain of GCN2 ^17–19^ inducing conformational changes ^20–21^ and autophosphorylation. Since hypoxia induces GCN2 phosphorylation at Thr899 in the kinase domain indicative of GCN2 activation in HLMVECs with complete growth medium as early as 2h exposure, hypoxia-induced activation of GCN2 is independent of the canonical uncharged tRNA binding, which is consistent with previous findings that GCN2 can be activated independent of tRNA binding, for example, mutations at or flanking the protein kinase domain of GCN2 cause constitutive activation.^25^ It is reported that GCN2 can be activated by hypoxia in mouse embryonic fibroblast cells,^28^ however, the potential mechanism leading to GCN2 activation is unclear. In our study, we have demonstrated that hypoxia activated and phosphorylated GCN2 through PDK1 not AMPK in ECs. We also demonstrated GCN2 activation in MCT-induced rat PAH model and in IPAH patients. Employing endothelium-targeted nanoparticles to deliver the CRISPR/Cas9 system^35,50^ to knockout *Gcn2* selectively in ECs, we also observed attenuated PH in these EC-specific knockout mice, demonstrating the important role of hypoxia-activated GCN2 in ECs in mediating pulmonary vascular remodeling and pulmonary hypertension development. Since we also observed GCN2 hyperphosphorylation in pulmonary vascular SMCs and interstitial tissues in MCT rats and IPAH patients, it is possible that GCN2 can also be activated by other PAH-causing factors such as growth factors or inflammatory mediators^51^ in other vascular cell types to mediate pulmonary vascular remodeling and PAH development. Future studies are warranted to delineate the molecular mechanisms leading to GCN2 activation in different cell types in response to various PAH-causing factors.

EDN1, an EC-derived factor, plays an important role in the pathogenesis of PAH in increasing vasoconstriction and cell proliferation. EDN1 exerts vasoconstrictor and mitogenic effects by binding to the two distinct receptors on vascular SMCs, making it the main therapeutic target in the treatment of PAH.^41–43^ Our study for the first time demonstrates the effect of GCN2 in regulating EDN1 expression in both the hypoxia PH mouse model and in cultured HLMVECs in response to hypoxia. *HIF1A* and *HIF2A* plasmid overexpression study demonstrated that EDN1 expression level inhibited by GCN2 deficiency was completely rescued by HIF-2a overexpression while only partially rescued by HIF-1a overexpression when same amount of HIF plasmid DNA was transfected to HLMVECs, revealing that GCN2 regulates EDN1 mainly through HIF-2a rather than HIF-1a as previously reported.^44,45^ Recent studies also support the role of HIF-2α in mediating EDN1 expression in mouse lungs and HLMVECs.^52,53^ Furthermore, we employed the endothelium-targeted nanoparticles to deliver *Edn1* gene specifically in ECs in KO mice and observed reversed PH phenotype in these mutant mice, demonstrating the casual role of GCN2/HIF-2α/EDN1 signaling axis in the mechanisms of pulmonary vascular remodeling and PH development in response to chronic hypoxia.

In addition to demonstrating the role of GCN2 activation in hypoxia-induced PH in mice, we also for the first time investigated the role of GCN2 in MCT-induced PAH in rats. GCN2 was found highly phosphorylated and activated in pulmonary vascular ECs of MCT rats. With compound A-92 treatment, MCT-induced PAH and pulmonary vascular remodeling were both suppressed, revealing the therapeutic potential of GCN2 inhibition for the treatment of PAH.

In conclusion, our data have demonstrated that disruption of the GCN2 kinase domain did not induce PVOD in mice, but unexpectedly inhibited pulmonary vascular remodeling and PH in response to chronic hypoxia. We found that hypoxia-induced Edn1 expression is mediated by GCN2 activation via HIF-2a expression, and that restored endothelial Edn1 expression in *Eif2ak4^-/-^*mice reversed the reduced PH phenotype in response to hypoxia, demonstrating the novel role of GCN2/HIF-2a/EDN1 signaling axis in the pathogenesis of pulmonary vascular remodeling and PAH. EC-specific disruption of *Eif2ak4* also attenuated hypoxia-induced PH in mice. Furthermore, MCT-induced PAH and vascular remodeling was suppressed by inhibiting GCN2 activity. Importantly, we also observed extensive GCN2 phosphorylation/activation in ECs of pulmonary vascular lesions of IPAH patients without marked changes of protein levels. Thus, targeting GCN2 signaling represents a novel therapeutic strategy for treatment of PAH patients, especially those without GCN2 loss-of-function mutations.

## Non-standard Abbreviations and Acronyms

ECs: endothelial cells
CRISPR: Clustered regularly interspaced short repeats, gRNA, guide RNA
HLMVECs: Human lung microvascular endothelial cells
LV+S: left ventricle+septum
MCT: monocrotaline
(I)PAH: (idiopathic) pulmonary arterial hypertension
PH: pulmonary hypertension
PVOD: pulmonary veno-occlusive disease
RV: right ventricle; RVSP, right ventricle systolic pressure
SMC: smooth muscle cells

## Acknowledgements

M.M.Z. and Y.Y.Z. conceived the experiments. M.M..Z, J.D, Z.D, Y.P designed, carried out experiments, and analyzed the data. Y.Y.Z. analyzed and interpreted the data. M.M.Z. draft the manuscript. Y.Y.Z. supervised the project and revised the manuscript and is responsible for the concept. All authors edited the manuscript.

## Source of Funding

This work was supported in part by NIH grants R01HL133951, R01HL140409, R01HL148810, R01HL164014, and R01HL162299 to Y.Y.Z. AHA predoctoral fellowship 19PRE34450171 to M.M.Z.

## Disclosures

Y.Y.Z is the founder and chief scientific officer of Mountview Therapeutics LLC. This project utilizes technologies subject to the following pending patent PCT/US2019/055787 “PLGA-PEG nanoparticles and methods of uses” by Zhao, Y.Y. The other authors declare no competing interests.

## Novelty and Significance

### What is known?

- Recessive mutations in the *EIF2AK4* gene (encoding GCN2) are reported in pulmonary veno-occlusive disease (PVOD) patients and pulmonary arterial hypertension (PAH) patients.
- GCN2 can be activated by amino acid starvation or hypoxia.

### What new information does this article contribute?

- *Eif2ak4* knockout mice with disruption of the kinase domain have neither spontaneous PVOD nor pulmonary hypertension (PH) but have reduced hypoxia-induced PH.
- Hypoxia activates PDK1/GCN2/HIF-2α/Edn1 axis in ECs, which plays a causal role in pulmonary vascular remodeling and PH development.
- Pharmacological inhibition of GCN2 inhibits PH in monocrotaline-challenged rats.

Our study has demonstrated that loss of GCN2 kinase function does not induce spontaneous PVOD and PH in mice but unexpectedly inhibits pulmonary vascular remodeling and PH development in response to chronic hypoxia. Pharmacological inhibition of GCN2 kinase activity also inhibits PAH in monocrotaline-challenged rats. GCN2 is markedly phosphorylated/activated in endothelial cells of both PAH patients and experimental PH rodents. Mechanistically, we have demonstrated that hypoxia activates GCN2 via PDK1 signaling and thus induces endothelin-1 expression through HIF-2α. In vivo rescue study shows that GCN2-regulated endothelin-1 expression plays a causal role in hypoxia-induced pulmonary vascular remodeling and PH development. Our study for the first time demonstrates a detrimental role of GCN2 activation in promoting pulmonary vascular remodeling and PAH development. Thus, targeting GCN2 signaling may represent a novel therapeutic approach for treatment of PAH patients without *EIF2AK4* mutations.

## References

1. Simonneau G and Hoeper MM. The revised definition of pulmonary hypertension: exploring the impact on patient management. Eur Heart J Suppl. 2019;21:K4–K8.

2. Hoeper MM, Humbert M, Souza R, Idrees M, Kawut SM, Sliwa-Hahnle K, Jing ZC and Gibbs JS. A global view of pulmonary hypertension. Lancet Respir Med. 2016;4:306–22.

3. Humbert M, Sitbon O and Simonneau G. Treatment of pulmonary arterial hypertension. N Engl J Med. 2004;351:1425–36.

4. McLaughlin VV, Archer SL, Badesch DB, Barst RJ, Farber HW, Lindner JR, Mathier MA, McGoon MD, Park MH, Rosenson RS, Rubin LJ, Tapson VF, Varga J, Harrington RA, Anderson JL, Bates ER, Bridges CR, Eisenberg MJ, Ferrari VA, Grines CL, Hlatky MA, Jacobs AK, Kaul S, Lichtenberg RC, Lindner JR, Moliterno DJ, Mukherjee D, Pohost GM, Rosenson RS, Schofield RS, Shubrooks SJ, Stein JH, Tracy CM, Weitz HH, Wesley DJ and Accf/Aha. ACCF/AHA 2009 expert consensus document on pulmonary hypertension: a report of the American College of Cardiology Foundation Task Force on Expert Consensus Documents and the American Heart Association: developed in collaboration with the American College of Chest Physicians, American Thoracic Society, Inc., and the Pulmonary Hypertension Association. Circulation. 2009;119:2250–94.

5. Galie N, Humbert M, Vachiery JL, Gibbs S, Lang I, Torbicki A, Simonneau G, Peacock A, Vonk Noordegraaf A, Beghetti M, Ghofrani A, Gomez Sanchez MA, Hansmann G, Klepetko W, Lancellotti P, Matucci M, McDonagh T, Pierard LA, Trindade PT, Zompatori M, Hoeper M and Group ESCSD. 2015 ESC/ERS Guidelines for the diagnosis and treatment of pulmonary hypertension: The Joint Task Force for the Diagnosis and Treatment of Pulmonary Hypertension of the European Society of Cardiology (ESC) and the European Respiratory Society (ERS): Endorsed by: Association for European Paediatric and Congenital Cardiology (AEPC), International Society for Heart and Lung Transplantation (ISHLT). Eur Heart J. 2016;37:67–119.

6. D’Alonzo GE, Barst RJ, Ayres SM, Bergofsky EH, Brundage BH, Detre KM, Fishman AP, Goldring RM, Groves BM, Kernis JT and, et al. Survival in patients with primary pulmonary hypertension. Results from a national prospective registry. Ann Intern Med. 1991;115:343–9.

7. Galie N, Barbera JA, Frost AE, Ghofrani HA, Hoeper MM, McLaughlin VV, Peacock AJ, Simonneau G, Vachiery JL, Grunig E, Oudiz RJ, Vonk-Noordegraaf A, White RJ, Blair C, Gillies H, Miller KL, Harris JH, Langley J, Rubin LJ and Investigators A. Initial Use of Ambrisentan plus Tadalafil in Pulmonary Arterial Hypertension. N Engl J Med. 2015;373:834–44.

8. Benza RL, Miller DP, Barst RJ, Badesch DB, Frost AE and McGoon MD. An evaluation of long-term survival from time of diagnosis in pulmonary arterial hypertension from the REVEAL Registry. Chest. 2012;142:448–456.

9. Sitbon O, Sattler C, Bertoletti L, Savale L, Cottin V, Jais X, De Groote P, Chaouat A, Chabannes C, Bergot E, Bouvaist H, Dauphin C, Bourdin A, Bauer F, Montani D, Humbert M and Simonneau G. Initial dual oral combination therapy in pulmonary arterial hypertension. Eur Respir J. 2016;47:1727–36.

10. van de Veerdonk MC, Kind T, Marcus JT, Mauritz GJ, Heymans MW, Bogaard HJ, Boonstra A, Marques KM, Westerhof N and Vonk-Noordegraaf A. Progressive right ventricular dysfunction in patients with pulmonary arterial hypertension responding to therapy. J Am Coll Cardiol. 2011;58:2511–9.

11. Lau EMT, Giannoulatou E, Celermajer DS and Humbert M. Epidemiology and treatment of pulmonary arterial hypertension. Nat Rev Cardiol. 2017;14:603–614.

12. Berlanga JJ, Santoyo J and De Haro C. Characterization of a mammalian homolog of the GCN2 eukaryotic initiation factor 2alpha kinase. Eur J Biochem. 1999;265:754–62.

13. Sood R, Porter AC, Olsen DA, Cavener DR and Wek RC. A mammalian homologue of GCN2 protein kinase important for translational control by phosphorylation of eukaryotic initiation factor-2alpha. Genetics. 2000;154:787–801.

14. Harding HP, Novoa I, Zhang Y, Zeng H, Wek R, Schapira M and Ron D. Regulated translation initiation controls stress-induced gene expression in mammalian cells. Mol Cell. 2000;6:1099–108.

15. Yang R, Wek SA and Wek RC. Glucose limitation induces GCN4 translation by activation of Gcn2 protein kinase. Mol Cell Biol. 2000;20:2706–17.

16. Rolfes RJ and Hinnebusch AG. Translation of the yeast transcriptional activator GCN4 is stimulated by purine limitation: implications for activation of the protein kinase GCN2. Mol Cell Biol. 1993;13:5099–111.

17. Dong J, Qiu H, Garcia-Barrio M, Anderson J and Hinnebusch AG. Uncharged tRNA activates GCN2 by displacing the protein kinase moiety from a bipartite tRNA-binding domain. Mol Cell. 2000;6:269–79.

18. Ruff M, Krishnaswamy S, Boeglin M, Poterszman A, Mitschler A, Podjarny A, Rees B, Thierry JC and Moras D. Class II aminoacyl transfer RNA synthetases: crystal structure of yeast aspartyl-tRNA synthetase complexed with tRNA(Asp). Science. 1991;252:1682–9.

19. Wek SA, Zhu S and Wek RC. The histidyl-tRNA synthetase-related sequence in the eIF-2 alpha protein kinase GCN2 interacts with tRNA and is required for activation in response to starvation for different amino acids. Mol Cell Biol. 1995;15:4497–506.

20. Qiu H, Garcia-Barrio MT and Hinnebusch AG. Dimerization by translation initiation factor 2 kinase GCN2 is mediated by interactions in the C-terminal ribosome-binding region and the protein kinase domain. Mol Cell Biol. 1998;18:2697–711.

21. Qiu H, Dong J, Hu C, Francklyn CS and Hinnebusch AG. The tRNA-binding moiety in GCN2 contains a dimerization domain that interacts with the kinase domain and is required for tRNA binding and kinase activation. EMBO J. 2001;20:1425–38.

22. Romano PR, Garcia-Barrio MT, Zhang X, Wang Q, Taylor DR, Zhang F, Herring C, Mathews MB, Qin J and Hinnebusch AG. Autophosphorylation in the activation loop is required for full kinase activity in vivo of human and yeast eukaryotic initiation factor 2alpha kinases PKR and GCN2. Mol Cell Biol. 1998;18:2282–97.

23. Wek RC, Cannon JF, Dever TE and Hinnebusch AG. Truncated protein phosphatase GLC7 restores translational activation of GCN4 expression in yeast mutants defective for the eIF-2 alpha kinase GCN2. Mol Cell Biol. 1992;12:5700–10.

24. Padyana AK, Qiu H, Roll-Mecak A, Hinnebusch AG and Burley SK. Structural basis for autoinhibition and mutational activation of eukaryotic initiation factor 2alpha protein kinase GCN2. J Biol Chem. 2005;280:29289–99.

25. Qiu H, Hu C, Dong J and Hinnebusch AG. Mutations that bypass tRNA binding activate the intrinsically defective kinase domain in GCN2. Genes Dev. 2002;16:1271–80.

26. Deng J, Harding HP, Raught B, Gingras AC, Berlanga JJ, Scheuner D, Kaufman RJ, Ron D and Sonenberg N. Activation of GCN2 in UV-irradiated cells inhibits translation. Curr Biol. 2002;12:1279–86.

27. Zhan K, Narasimhan J and Wek RC. Differential activation of eIF2 kinases in response to cellular stresses in Schizosaccharomyces pombe. Genetics. 2004;168:1867–75.

28. Liu Y, Laszlo C, Liu Y, Liu W, Chen X, Evans SC and Wu S. Regulation of G(1) arrest and apoptosis in hypoxia by PERK and GCN2-mediated eIF2alpha phosphorylation. Neoplasia. 2010;12:61–8.

29. Schmidt S, Gay D, Uthe FW, Denk S, Paauwe M, Matthes N, Diefenbacher ME, Bryson S, Warrander FC, Erhard F, Ade CP, Baluapuri A, Walz S, Jackstadt R, Ford C, Vlachogiannis G, Valeri N, Otto C, Schulein-Volk C, Maurus K, Schmitz W, Knight JRP, Wolf E, Strathdee D, Schulze A, Germer CT, Rosenwald A, Sansom OJ, Eilers M and Wiegering A. A MYC-GCN2-eIF2alpha negative feedback loop limits protein synthesis to prevent MYC-dependent apoptosis in colorectal cancer. Nat Cell Biol. 2019;21:1413–1424.

30. Ravindran R, Loebbermann J, Nakaya HI, Khan N, Ma H, Gama L, Machiah DK, Lawson B, Hakimpour P, Wang YC, Li S, Sharma P, Kaufman RJ, Martinez J and Pulendran B. The amino acid sensor GCN2 controls gut inflammation by inhibiting inflammasome activation. Nature. 2016;531:523–527.

31. Eyries M, Montani D, Girerd B, Perret C, Leroy A, Lonjou C, Chelghoum N, Coulet F, Bonnet D, Dorfmuller P, Fadel E, Sitbon O, Simonneau G, Tregouet DA, Humbert M and Soubrier F. EIF2AK4 mutations cause pulmonary veno-occlusive disease, a recessive form of pulmonary hypertension. Nat Genet. 2014;46:65–9.

32. Montani D, Price LC, Dorfmuller P, Achouh L, Jais X, Yaici A, Sitbon O, Musset D, Simonneau G and Humbert M. Pulmonary veno-occlusive disease. Eur Respir J. 2009;33:189–200.

33. Hadinnapola C, Bleda M, Haimel M, Screaton N, Swift A, Dorfmuller P, Preston SD, Southwood M, Hernandez-Sanchez J, Martin J, Treacy C, Yates K, Bogaard H, Church C, Coghlan G, Condliffe R, Corris PA, Gibbs S, Girerd B, Holden S, Humbert M, Kiely DG, Lawrie A, Machado R, MacKenzie Ross R, Moledina S, Montani D, Newnham M, Peacock A, Pepke-Zaba J, Rayner-Matthews P, Shamardina O, Soubrier F, Southgate L, Suntharalingam J, Toshner M, Trembath R, Vonk Noordegraaf A, Wilkins MR, Wort SJ, Wharton J, Consortium NB-RD, Idiopathic UKNCSo, Heritable PAH, Graf S and Morrell NW. Phenotypic Characterization of EIF2AK4 Mutation Carriers in a Large Cohort of Patients Diagnosed Clinically With Pulmonary Arterial Hypertension. Circulation. 2017;136:2022–2033.

34. Eichstaedt CA, Song J, Benjamin N, Harutyunova S, Fischer C, Grunig E and Hinderhofer K. EIF2AK4 mutation as “second hit” in hereditary pulmonary arterial hypertension. Respir Res. 2016;17:141.

35. Zhang X, Jin H, Huang X, Chaurasiya B, Dong D, Shanley TP and Zhao YY. Robust genome editing in adult vascualr ensdothelium by nanoparticle delivery of CRISPR-Cas9 plasmid DNA. Cell Rep. 2022;38:110196.

36. Hamanaka RB, Bennett BS, Cullinan SB and Diehl JA. PERK and GCN2 contribute to eIF2alpha phosphorylation and cell cycle arrest after activation of the unfolded protein response pathway. Mol Biol Cell. 2005;16:5493–501.

37. Cherkasova V, Qiu H and Hinnebusch AG. Snf1 promotes phosphorylation of the alpha subunit of eukaryotic translation initiation factor 2 by activating Gcn2 and inhibiting phosphatases Glc7 and Sit4. Mol Cell Biol. 2010;30:2862–73.

38. Kimpe M, Voordeckers K, Thevelein JM and Van Zeebroeck G. Pkh1 interacts with and phosphorylates components of the yeast Gcn2/eIF2alpha system. Biochem Biophys Res Commun. 2012;419:89–94.

39. Najafov A, Sommer EM, Axten JM, Deyoung MP and Alessi DR. Characterization of GSK2334470, a novel and highly specific inhibitor of PDK1. Biochem J. 2011;433:357–69.

40. Kourembanas S, Marsden PA, McQuillan LP and Faller DV. Hypoxia induces endothelin gene expression and secretion in cultured human endothelium. J Clin Invest. 1991;88:1054–7.

41. Prie S, Leung TK, Cernacek P, Ryan JW and Dupuis J. The orally active ET(A) receptor antagonist (+)-(S)-2-(4,6-dimethoxy-pyrimidin-2-yloxy)-3-methoxy-3,3-diphe nyl-propionic acid (LU 135252) prevents the development of pulmonary hypertension and endothelial metabolic dysfunction in monocrotaline-treated rats. J Pharmacol Exp Ther. 1997;282:1312–8.

42. Satwiko MG, Ikeda K, Nakayama K, Yagi K, Hocher B, Hirata K and Emoto N. Targeted activation of endothelin-1 exacerbates hypoxia-induced pulmonary hypertension. Biochem Biophys Res Commun. 2015;465:356–62.

43. Giaid A, Yanagisawa M, Langleben D, Michel RP, Levy R, Shennib H, Kimura S, Masaki T, Duguid WP and Stewart DJ. Expression of endothelin-1 in the lungs of patients with pulmonary hypertension. N Engl J Med. 1993;328:1732–9.

44. Yamashita K, Discher DJ, Hu J, Bishopric NH and Webster KA. Molecular regulation of the endothelin-1 gene by hypoxia. Contributions of hypoxia-inducible factor-1, activator protein-1, GATA-2, AND p300/CBP. J Biol Chem. 2001;276:12645–53.

45. Minchenko A and Caro J. Regulation of endothelin-1 gene expression in human microvascular endothelial cells by hypoxia and cobalt: Role of hypoxia responsive element. Mol Cell Biochem 2000;208:53–62.

46. Brazeau JF and Rosse G. Triazolo[4,5-d]pyrimidine Derivatives as Inhibitors of GCN2. ACS Med Chem Lett. 2014;5:282–3.

47. Emanuelli G, Nassehzadeh-Tabriz N, Morrell NW and Marciniak SJ. The integrated stress response in pulmonary disease. Eur Respir Rev. 2020;29.

48. Nossent EJ, Antigny F, Montani D, Bogaard HJ, Ghigna MR, Lambert M, Thomas de Montpreville V, Girerd B, Jais X, Savale L, Mercier O, Fadel E, Soubrier F, Sitbon O, Simonneau G, Vonk Noordegraaf A, Humbert M, Perros F and Dorfmuller P. Pulmonary vascular remodeling patterns and expression of general control nonderepressible 2 (GCN2) in pulmonary veno-occlusive disease. J Heart Lung Transplant. 2018;37:647–655.

49. Lai YJ, Chen PR, Huang YL and Hsu HH. Unique wreath-like smooth muscle proliferation of the pulmonary vasculature in pulmonary veno-occlusive disease versus pulmonary arterial hypertension. J Formos Med Assoc. 2020;119:300–309.

50. Huang X, Zhang X, Machireddy N, Evans CE, Trewartha SD, Hu G, Mutlu GM, Fang Y, Wu D, and Zhao YY. Endothelial FoxM1 reactivates aging-impaired endothelial regeneration for vascular repair and resolution of inflammatory lung injury. Sci Transl Med. 2023;15:eabm5755.

51. Evans CE, Cober ND, Dai Z, Stewart DJ and Zhao YY. Endothelial cells in the pathogenesis of pulmonary arterial hypertension. Eur Respir J. 2021;58.

52. Dai Z, Zhu MM, Peng Y, Jin H, Machireddy N, Qian Z, Zhang X and Zhao YY. Endothelial and Smooth Muscle Cell Interaction via FoxM1 Signaling Mediates Vascular Remodeling and Pulmonary Hypertension. Am J Respir Crit Care Med. 2018;198:788–802.

53. Dai Z, Li M, Wharton J, Zhu MM, and Zhao YY. Prolyl-4 hydroxylase 2 (PHD2) deficiency in endothelial cells and hematopoietic cells induces obliterative vascular remodeling and severe pulmonary arterial hypertension in mice and humans through hypoxia-inducible factor-2α. Circulation. 2016;133:2447–2458.

